# Serpins: Purification and characterization of potent protease inhibitors from *Clostridium thermocellum*

**DOI:** 10.1101/2020.04.21.053413

**Authors:** Maria Mushtaq, Muhammad Javaid Asad, Muhammad Zeeshan Hyder, Syed Muhammad Saqlan Naqvi, Saad Imran Malik, Raja Tahir Mehmood

## Abstract

*Clostridium thermocellum* produces an extracellular cellulosome (a multiprotein complex produced by firmicutes bacteria), which, owing to its extracellular location, is open to protease attack. Serine protease inhibitors (serpins) protect bacteria against protease attack. However, their structure and function are poorly characterized. This study identified and amplified the serpin 1270 gene from the C. thermocellum genome. Purified serpins were cloned into the pTXB1 vector using the one-step sequence and ligation-independent cloning reaction and transformed into Escherichia coli BL21 DE3 cells. Enzyme overexpression and purification and enzyme inhibitory assays were performed. The results showed that serpin 1270 has 89% inhibition against Bacillus subtilisin and 64% inhibition against trypsin, chymotrypsin, and papain.

## Introduction

Serine protease inhibitors (serpins) are a major, largely distributed superfamily of protease inhibitors, recognized half a century ago. There are 68 classes of protease inhibitors, and serpins are the largest class; protein sequences are available in the SWISS-PROT and TrEMBL databases (1). More than 1500 serpin genes have been identified in animals, plants, and prokaryotes. In multicellular eukaryotes, serpin genes have been identified in humans, *Drosophila*, *Arabidopsis thaliana*, and *Caenorhabditis elegans*. In prokaryotes, their distribution is irregular, in single-gene form. Serine proteases are the major target of serpins, but some serpins, such as cysteine proteases (e.g., viral serpin crmA with caspase-1, SCCA-1, and MENT), inhibit proteases other than serine proteases (2). Not all serpins are inhibitory proteins; some noninhibitory serpins play a major role in physiological pathways such as blood coagulation, fibrinolysis, hormone binding, intracellular signaling, and programmed cell death and can be used as a therapeutic agent against gram-positive and gram-negative bacteria (3,4,5). Maspins, a class of mammary serpins, can be used as a potential marker for the screening of esophageal cancers (6). Serpins have a highly conserved structure, comprising 300–500 amino acids with 3 β-sheets and 7–9 α-helices. A stretch of ~17 preserved amino acids is present in the middle of β-sheets A and C, which act as protein recognition sites. This stretch is called the reactive center loop. Bioinformatics analysis has identified serpins in the animal and plant kingdoms, and it is believed that they have common evolutionary ancestors (fungi and prokaryotes). However, no fungal serpin has been identified, and little is known about prokaryotic serpins as well (7,8)

These findings pave the way for the identification of prokaryotic serpins and their possible functions. In an effort to find serpins in prokaryotes, two serpin genes were identified in an anaerobic gram-positive bacterium, *Clostridium thermocellum* (9). *C. thermocellum* is the most investigated anaerobic bacterium because of its ability to degrade cellulose and directly convert it into ethanol. It is the most efficient biomass degrader, and because of its recent rise in popularity, it is well characterized. It can form biofilm, where it arranges itself in a parallel position, and it can grow on different substrates besides cellulose (e.g., cellobiose, xylose, and hemicellulose). Its growth temperature range is 50°C–68°C, which is suitable for industrial processes (10). Despite features that make *C. thermocellum* an ideal candidate for fermentation, its most important characteristic is the presence of cellulosome, a discrete enzyme unit and a multienzyme machinery comprising 20 different enzymes that aid in cellulose degradation (11). In addition, cellulosome contains serpins.

The primary function of serpins is to neutralize the effect of serine proteases (12). Kang et al. (4), for the first time, reported the presence of two serpin-like genes in the *C. thermocellum* cellulosome, which showed high sequence identity with the serpin from *Thermobifida fusca*, which also contained a dockerin module like those of other dockerins present in the *C. thermocellum* cellulosome.

*C. thermocellum* is a gram-positive anaerobic bacterium that limits its culturing, and serpin 1270 is an extracellulosmal enzyme that can affect the activity of cellulases. The structure and function of serpins are poorly characterized. This study aimed to identify the serpins present in prokaryotes. Keeping in view the importance of serpin genes and their presence in the *C. thermocellum* genome, we cloned, purified, and characterized the *serpin 1270* gene from the *C. thermocellum* genome.

## Materials and Methods

### Bacterial strains

The genomic DNA of *C. thermocellum* was obtained from the Department of Chemical Engineering and Applied Sciences, University of Rochester, USA. *Escherichia coli* BL21 DE3 cells were used for the cloning and expression of *C. thermocellum serpin 1270*.

### Amplification of *serpin 1270*

#### Primer designing and polymerase chain reaction

Sequences of *serpin 1270* were obtained from Artemis, and the complete genome sequence of *C. thermocellum* was screened for the presence of serpin genes. The following sequences of degenerate primers were designed manually to amplify *serpin 1270*: *serpin* 1270 forward primer: 5′ ATG AAA AGA ACA GCT TGC G 3′ *serpin 1270* reverse primer: 5′ ATA TTT TTC ACA ATC ATA CAA TTT G 3′

The restriction sites of the two enzymes, Nde-1 and Sap-1, were introduced at the 5′ end of the degenerate primers by introducing overhang regions, that is, forward (Nde-1: 5′ ACTTTAAGAAGGAGATATA<u>CATATG</u> 3′) and reverse (Sap-1: 5′ GGCAACTAGT<u>GCTCTC</u>CCGTGATGCA 3′), to clone *serpin 1270* into the pTXB1 vector under T7 promoter expression. The primers were analyzed using the oligo IDT analyzer available at http://idtdna.com/analyzer/applications/oligoanalyzer.

Polymerase chain reaction (PCR) conditions were optimized to perform PCR amplification of *C. thermocellum serpin 1270:* preamplification denaturation for 3 min at 94°C, denaturation for 30 s at 94°C, annealing (3 cycles) for 30 s at 50°C and extension for 2 min 15 s at 74°C, followed by 32 cycles of 30 s at 60°C and extension for 2 min 15 s at 74°C. The final extension was performed for 7 min at 74°C. The total reaction volume was 50 µL (Table 1). Then, 1% agarose gel electrophoresis was used to analyze the amplified products, and the fragments of interest were purified using the QIAGEN PCR purification kit (QIAGEN, Germany). The concentration of the amplified DNA fragments was monitored using a nanodrop spectrophotometer at Medical Center, University of Rochester.

**Table 1.** Composition of the reaction mixture for PCR amplification

| **Number.** | **Components** | **Concentration (uL)** |
| --- | --- | --- |
| 1. | 5× iProof HF buffer | 10 |
| 2. | 10 mM dNTPs | 1 |
| 3. | Forward Primer | 2.5 |
| 4. | Reverse Primer | 2.5 |
| 5. | Template DNA (50 ng/μL) | 5 |
| 6. | iProof DNA polymerase | 0.5 |
| 7. | Sterile H<sub>2</sub>O | 28.5 |

PCR: polymerase chain reaction

### Cloning and transformation of *serpin 1270*

#### Plasmid isolation and construction of digested plasmid

For the cloning and expression of serpins, the pTXB1 plasmid was isolated from *E. coli* XL-1 blue cells (Stratagene California, San Diego, CA, USA). For the cloning and ligation of the amplified gene, the pTXB1 plasmid was double-digested with Nde-I and Sap-I (New England Biolabs, Ipswich, MA, USA) enzymes at the respective restriction sites to generate binding sites for insert ligation. The total reaction volume was 30 μL, containing 20 μL of purified pTXB1 plasmid, 1.5 μL of Nde-1, 1.5 μL of Sap-1, and 3 μL of CutSmart buffer (New England Biolabs) with 4 μL of sterile water. Incubation was performed for 3 h in a hot water bath at 37°C. The digested pTXB1 plasmid was visualized using 1% agarose gel electrophoresis and purified using the QIAGEN gel extraction system according to the manufacturer’s instructions. The concentration of the purified double-digested pTXB1 plasmid was monitored using a nanodrop spectrophotometer.

#### Ligation reaction

The amplified products were ligated into the purified digested PTXB1 vector using the one-step sequence and ligation-independent cloning reaction with 150 ng of the plasmid and the PCR insert in a 1:4 molar ratio. The total reaction volume was 10 μL containing 2.5 μL of 150 ng of pTXB1, 1.5 μL of T4 DNA polymerase, and NEB2 buffer with 1 μL of bovine serum albumin (BSA), and the amplified product was incubated for 2.5 min at room temperature and then quenched on ice for 10 min for annealing.

#### Transformation in E. coli

For *serpin 1270* cloning and expression, *E. coli* BL21 DE3 cells were transformed with a ligation mixture of recombinant pTXB1 plasmid using the heat-shock method. Briefly, 1 μL of the reaction mixture was mixed with 70 μL of competent cells and kept on ice for 25 min. Then, a heat-shock was applied for 45 s at 42°C, followed by the addition of 1 mL of prewarmed lysogeny broth (LB) medium. The tubes were incubated for 1 h at 37°C with continuous shaking. Cells were pelleted down by mild centrifugation, the pellet was suspended in 200 μL of the medium, and the supernatant was discarded. Next, 100 μL of transformation mixture was spread on LB plates containing 100 mg/mL of ampicillin. Bacteria with transformed genes were selected on the basis of antibiotic resistance. Further confirmation for positive transformants was also carried out by colony PCR and restriction digestion of the recombinant plasmid.

The transformation efficiency was calculated as follows:

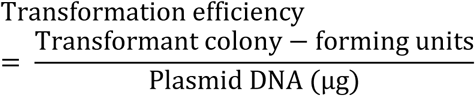

### *Serpin 1270* overexpression

#### Preparation of starter cultures

For *serpin 1270* expression, 3 mL of LB medium was inoculated with recombinant pTXB1 plasmid glycerol stocks. Cultures were incubated overnight at 250 rpm with recombinant pTXB1 by inoculation of positive clones from glycerol stocks. Then, 15 mL of fresh LB medium with the required antibiotic was added to 1 mL of bacterial cultures and incubated for 2–3 h at 37°C with continuous shaking until the optical density at a wavelength of 600 nm (OD_600_) reached the 0.5 early log phase of cells. The cells were induced with 1 mM isopropyl-β-D-thiogalactopyranoside (IPTG) and incubated for 15 h at 16°C and 250 rpm to achieve *serpin 1270* overexpression. The cells were harvested by centrifugation of the bacterial cultures at 6000 × *g* for 20 min. The supernatant was discarded, and pellets were stored at –80°C for future use. To check *serpin 1270* expression, induced and uninduced cells were resolved using 11% sodium dodecyl sulfate– polyacrylamide gel electrophoresis (SDS-PAGE) (13).

### Purification of recombinant *serpin 1270* and protein estimation

Expressed proteins were purified using the intein-mediated purification and affinity chitin-binding tag (IMPACT) chitin affinity matrix system (New England Biolabs). A cell pellet was dissolved in 8 mL of column buffer (20 mM Tris-HCl, pH 8.5; 500 mM NaCl; and 1 mM ethylenediaminetetraacetic acid). The column was packed with chitin beads of 2 mL bed volume and washed with 10× volumes of column buffer (2 × 8 mL) before loading the crude cell extract. Then, the column was loaded with the clarified cell extract dissolved in 8 mL of column buffer and run through the column five times at a flow rate of 1 mL/min, followed by washing with 20 bed volumes of column buffer (2 × 10 mL). Recombinant protein was eluted after 20 h of incubation of the chitin column with 50 mM cleavage buffer (1900 μL of column buffer with 100 μL of 1000 mM dithiothreitol), and the target protein was released from the intein tag in 1 mL microcentrifuge tubes. Elution was run using 11% SDS-PAGE for purification analysis. Chitin resins were regenerated by washing the column with 3 bed volumes of 0.3 M NaOH (stripping solution confirmation). A biuret assay was performed to determine the protein concentration in the crude enzyme sample at an absorbance wavelength of 540 nm against BSA as the standard.

#### Inhibitory assay for recombinant serpin

The caseinolytic proteinase activity assay was used to find changes in carboxypeptidase-, chymotrypsin-, trypsin-, papain-, and *Bacillus* subtilisin–mediated proteolysis by purified *C. thermocellum*. The inhibitory activity of serpin against *Bacillus* subtilisin (protease; Sigma-Aldrich, St. Louis, MO, USA) was determined according to Ref. (14) using casein as the substrate. Briefly, 0.01 mL of 1 mg/mL of subtilisin was incubated with 0.2 mL of 1% casein solution in 1 mL of reaction buffer (100 mM Tris-HCl, pH 8.0) for 30 min at 37°C. Casein was digested by subtilisin. After incubation, 1 mL of 20% trichloroacetic acid was added to precipitate unhydrolyzed casein by centrifugation for 5–10 min. The supernatant was removed to determine the digested protein concentration. The inhibitory activity of purified recombinant protein was determined by the preincubation of subtilisin with 0.075 mL of purified recombinant *serpin 1270* for 15 min at 37°C, and the reaction was carried out as described earlier. One unit of enzyme inhibition activity is defined as the amount of inhibitor that decreases protease activity to 50% of the original value. The difference in the absolute value between the negative and the positive control was taken as 100% inhibition by serpins. Tyrosine (Sigma-Aldrich) was used as the standard for the enzyme inhibitory assay. A standard curve was plotted against different concentrations of tyrosine on the *x* axis and absorbance on the *y* axis (Fig. 1).

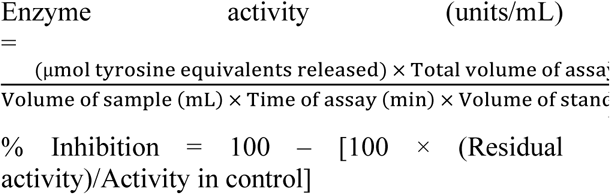

**Fig. 1.**
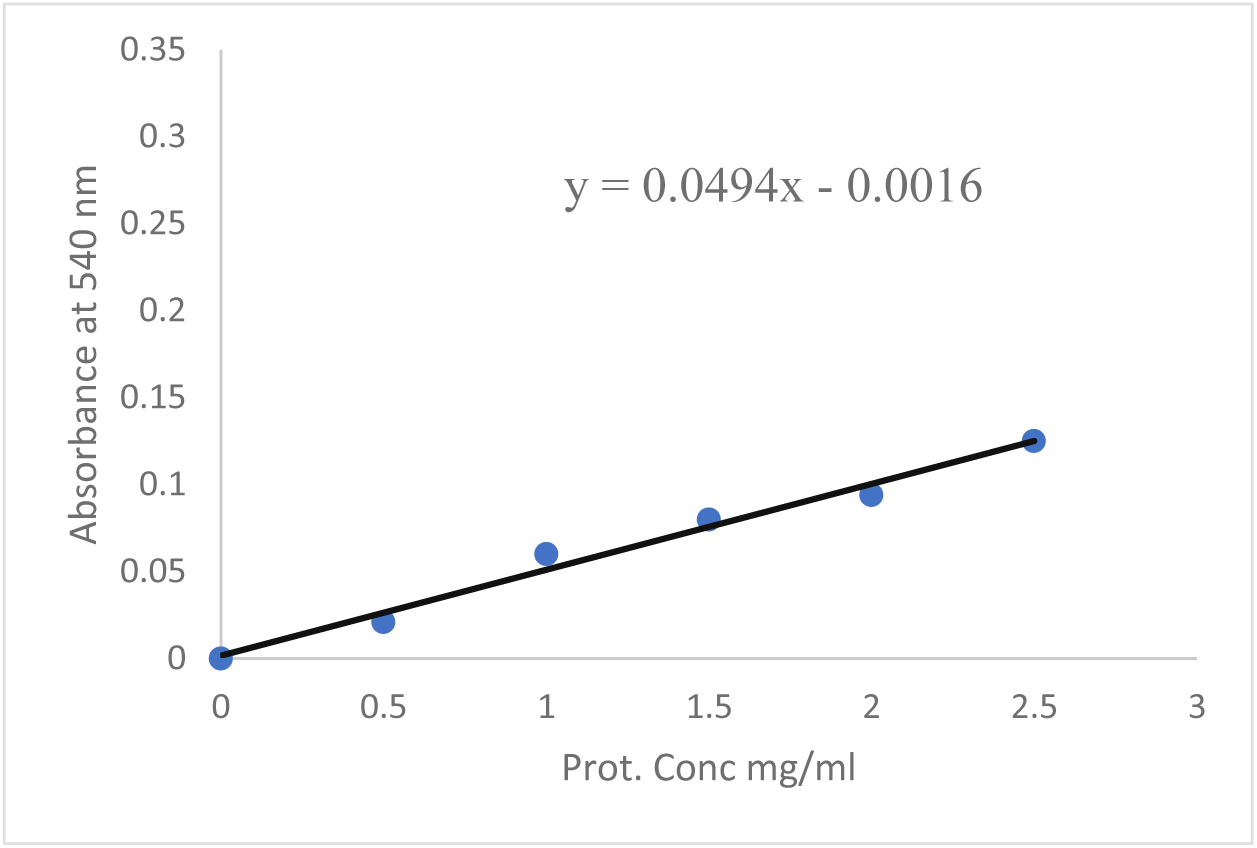
BSA standard curve at 540 nm. BSA: bovine serum albumin; PCR: polymerase chain reaction

### Statistical analysis

Enzyme activity was analyzed using analysis of variance (ANOVA) under a complete randomized design. The assay was performed in duplicate, and the mean with standard deviation was calculated for each treatment.

## Results and Discussion

### Screening, amplification, and purification of *C. thermocellum serpin 1270*

The gene sequence coding *serpin 1270* was identified with 1224 bp. The predicted relative molecular mass of *serpin 1270* was 44.8 kDa (Figs. 2 and 3).

**Fig. 2.**
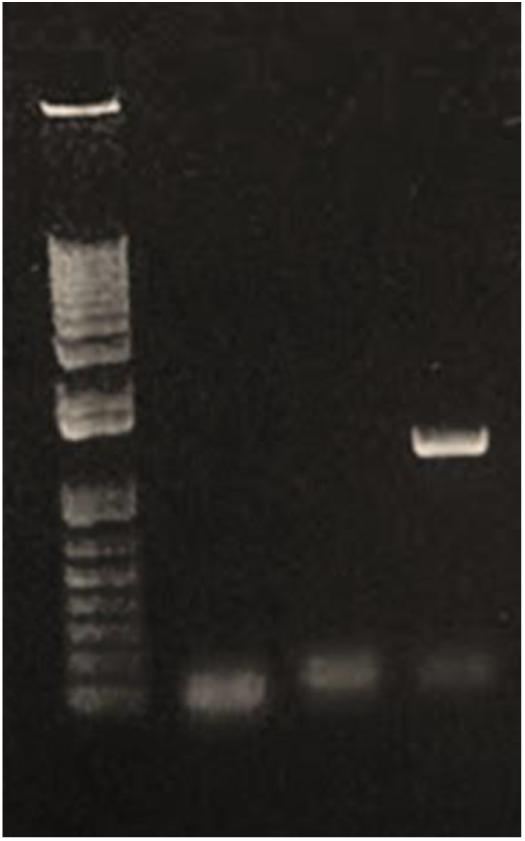
PCR amplification of *serpin 1270.* Amplified DNA products were analyzed in 1% agarose gel. Lane M: 1 Kb DNA ladder, gene *Cthe*_191 (1803 bp); lane 1: negative control (with no template); lane 3: amplified product of *serpin 1270*. The expected product of 1224 bp was obtained as a result of PCR amplification with degenerate primers on the *C. thermocellum* genomic DNA. PCR: polymerase chain reaction

**Fig. 3.**
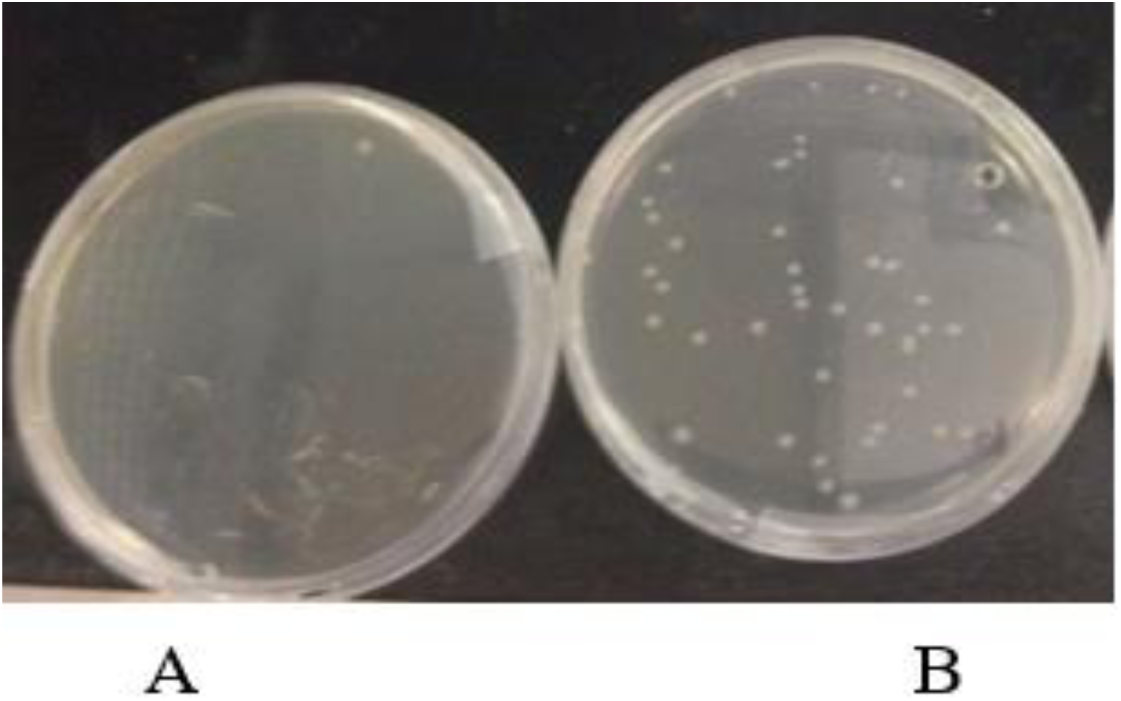
Transformation of recombinant *C. thermocellum serpin 1270* in *E. coli* BL21 DE3 cells. BL21 DE3 cells were transformed with cloned *serpin 1270* using the heat-shock method, and positive transformants were selected on the basis of ampicillin resistance against the control. A, negative control: cells without the pTXB1 plasmid; B: transformed cell with *Cthe_1270*.

### Cloning and overexpression of *serpin 1270*

The IMPACT system is a novel protein purification system that utilizes the self-cleavage of the target protein from the intein tag, which is an affinity tag with a chitin-binding domain for chitin resins. The pTXB1 plasmid allows fusion of the target protein at the C-terminus between the Nde1 and Sap1 sites without an extra amino acid on the target protein after cleavage of the intein tag. The pTXB1 plasmid is under the control of the T7 promoter, which is induced by IPTG, a lactose analog.

#### *Cloning of* serpin 1270

Positive transformants were selected on the basis of the ampicillin resistance gene (Fig. 4), and the pTXB1 plasmid was purified from the selected colonies. The cloning reaction was confirmed on 1% agarose gel electrophoresis (Fig. 5). The amplified product was obtained from 10 positive colonies, and transformation was confirmed (Fig. 6). The expected product of 8000 bp was obtained from samples 1, 2, 3, 4, 5, 7, 8, and 10. A positive colony was further confirmed by restriction digestion. The plasmid is digested by the enzymes (µmol tyrosine equivalents released) × Total volume of assayN(dmeL1) and Spe1 because there is a restriction in *serpin 1270*. Two fragments of 1249 and 6635 bp were obtained (Fig. 7).

**Fig. 4.**
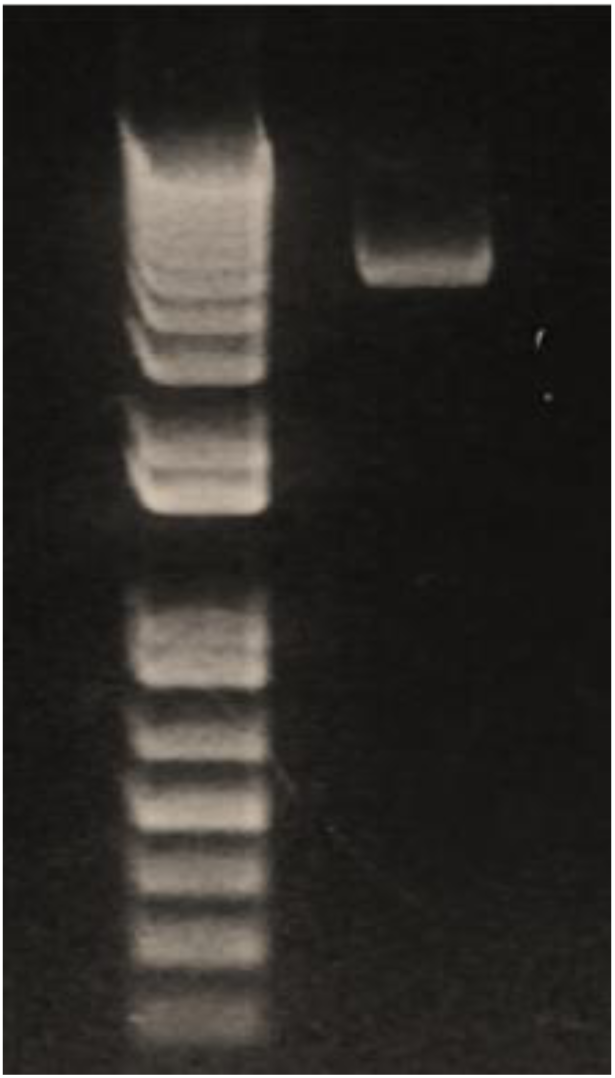
Confirmation of the cloning reaction by plasmid isolation from positive transformants. The pTXB1 plasmid was isolated from positive transformants (selected randomly) and analyzed on 1% agarose gel electrophoresis against the positive control. Lane M: molecular ladder (1 kb plus); lane 1: pTXB1 plasmid with insert *Cthe_1270* (7884 bp).

**Fig. 5.**
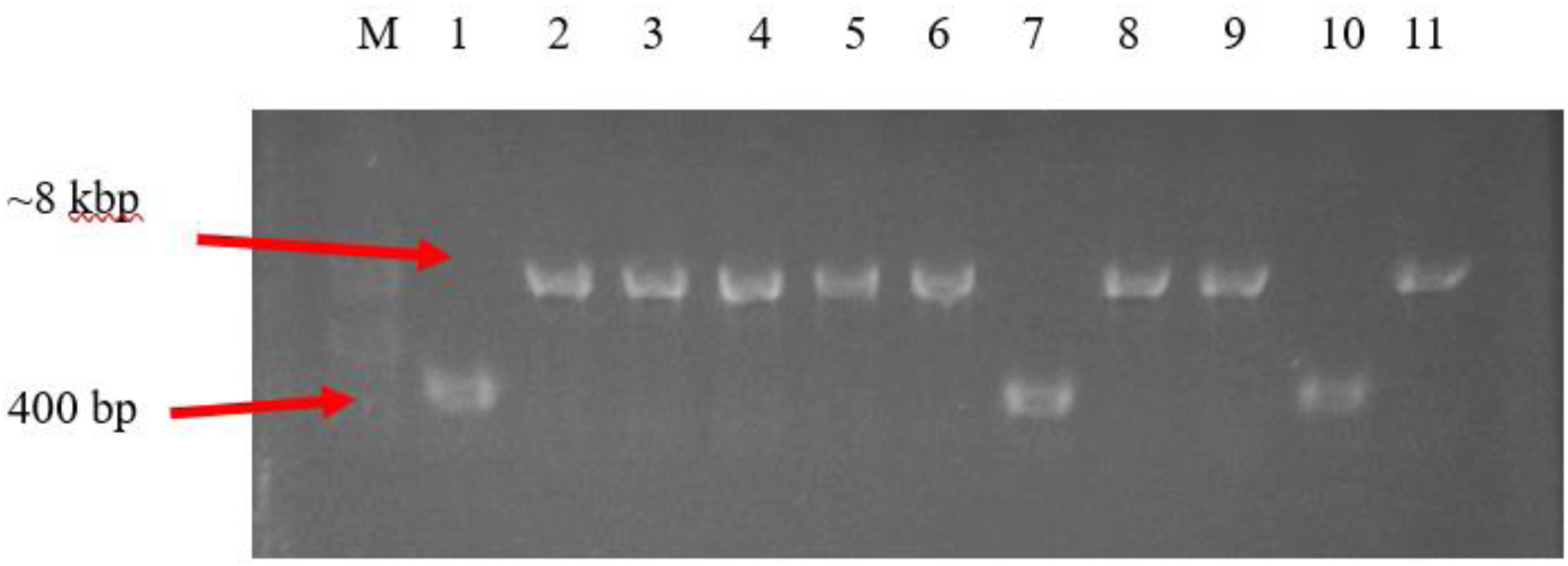
Colony PCR results obtained from transformed colonies of *E. coli* BL21 DE3 cells. Identification of *serpin 1270* from transformed colonies with degenerate primers pTXB1-F/-R. Lane M: 1 Kb DNA ladder; lane 1: positive control (pTXB1 with no gene); lanes 2–11: amplified product from transformed colonies 1–10. The expected product of 8000 bp was obtained from samples 1, 2, 3, 4, 5, 7, 8, and 10.

**Fig. 6.**
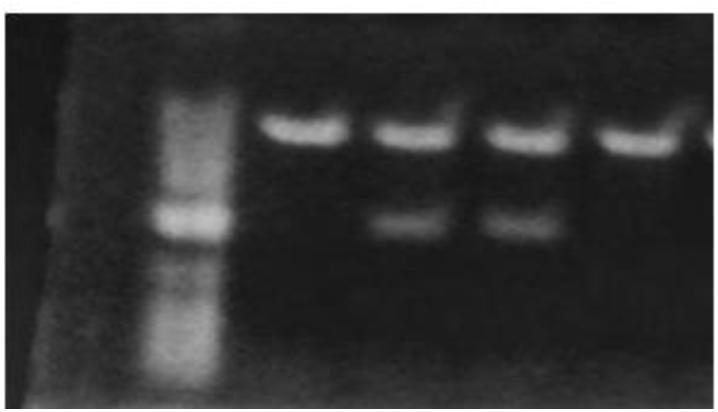
Confirmation of transformation by restriction digestion of the recombinant cloning vector. Digested and undigested recombinant pTXB1 cloning plasmid was analyzed on 1% agarose gel for further confirmation. Restriction digestion of the recombinant pTXB1 cloning plasmid was carried out with NdeI and SpeI. Lane M: 1 Kb DNA ladder; lane 1: negative control (pTXB1 plasmid with no gene); lanes 2 and 3: digested recombinant pTXB1 cloning plasmid with *serpin 1270* from samples 2 and 4. Expected products of 6.6 kb and 1.2 kbp were obtained. The 6.6 kb band shows a linearized pTXB1 cloning plasmid produced by releasing 1.2 kbp NdeI/SpeI-digested product.

**Fig. 7.**
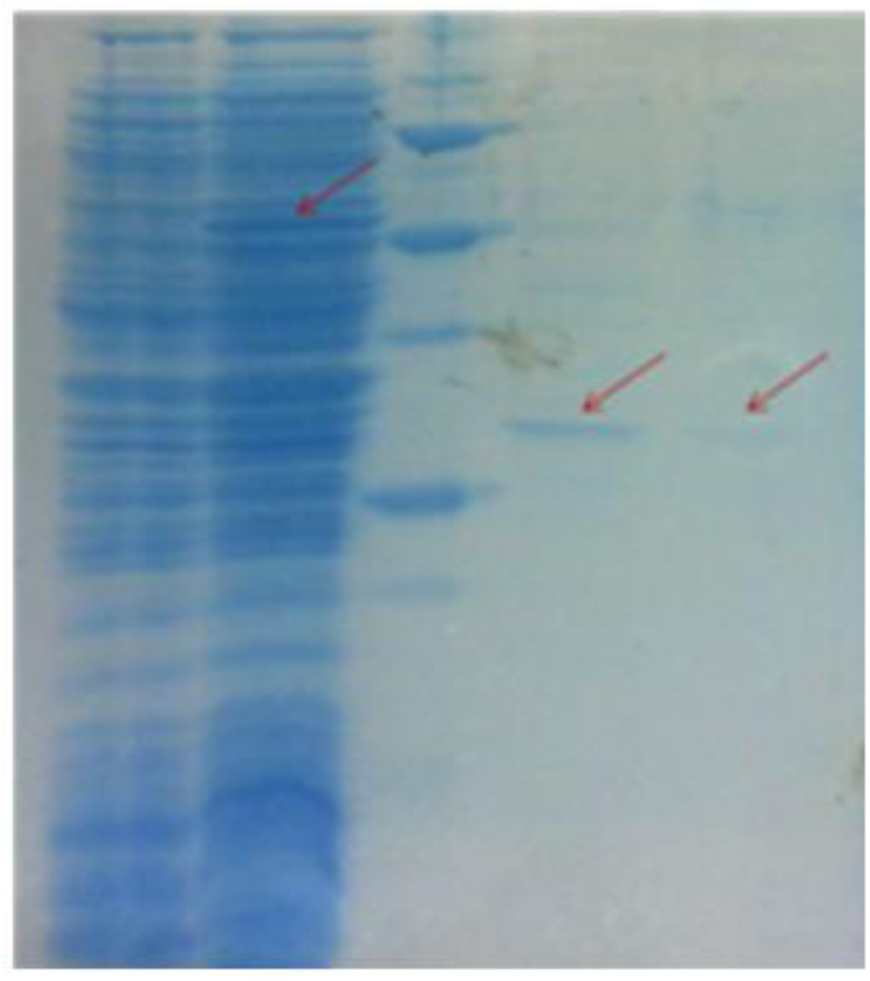
Expression and purification of *serpin 1270.* Lane 1: uninduced positive transformants *serpin 1270*; lane 2: induced positive transformants; lane M: protein molecular ladder; lane 3: elution 6 after purification; lane 4: elution 5 after purification.

#### *Expression levels of C.* thermocellum serpins 1270

Specific bands of 72 kDa for *serpin 1270* were observed on SDS-PAGE, containing a 28 kDa intein tag fused with overexpressed proteins (Fig. 8).

**Fig. 8.**
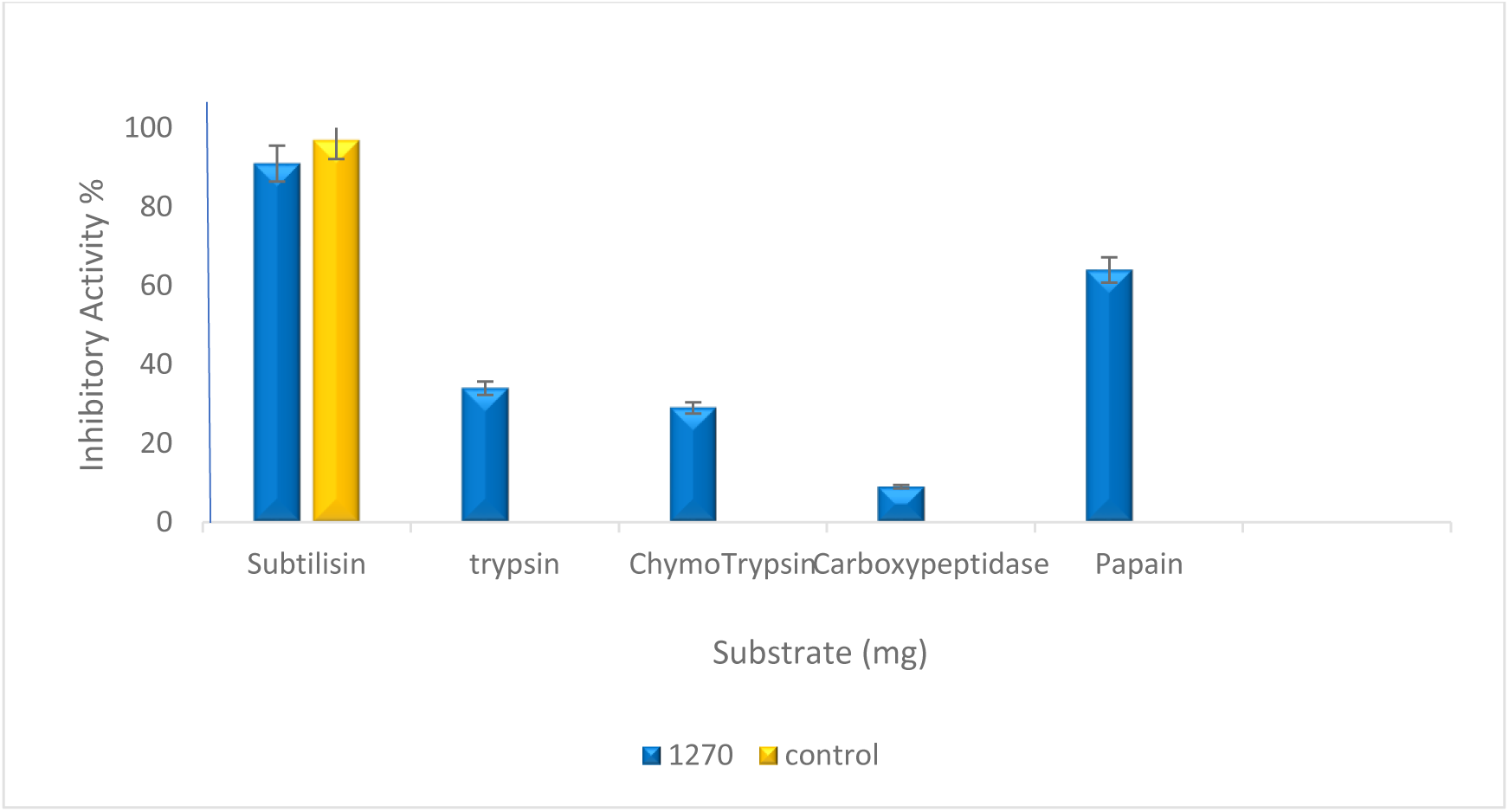
Inhibitory activity of *serpin 1270* against different proteases. *C. thermocellum serpin 1270* showed the maximum inhibitory activity of 89% against bacterial protease (subtilisin) and also showed 64% inhibitory activity against trypsin and chymotrypsin (mammalian proteases) and papain (a plant protease). *Bacillus subtilus* subtilase was used as a positive control. The data obtained were analyzed by ANOVA (*α* = 0.005). *P* < 0.01 and a coefficient of variation <10% indicated that the results are highly significant. ANOVA: analysis of variance

### Characterization of recombinant *serpins 1270*

#### Protein estimation and purification

The inhibitory activity against subtilisin after purification of crude enzyme increased, indicating that the *serpin 1270* concentration increases with purification. However, there was a decrease in the protein concentration due to removal of nonspecific proteins from the crude enzyme solution. Table 2 shows the purification values for recombinant serpins.

**Table 2.** Purification details for recombinant serpins from *C. thermocellum*

|  | **Volume**<br>**(ml)** | **%Inhib**<br>**ition** | **Protein**<br>**Concentration** | **Total**<br>**inhibition** | **Total**<br>**protein** | **Specific**<br>**activity** | **Purificatio**<br>**n fold** |
| --- | --- | --- | --- | --- | --- | --- | --- |
| **Crude**<br>**Enzyme** | 5 | 99 | 7.46 | 505 | 37.30 | 13.27 | 100 |
| **Serpin**<br>**1270** | 1 | 92 | 0.0912 | 92 | 0.0912 | 998.91 | 9.98 |

#### Substrate optimization

The maximum inhibition of purified *serpin 1270* against *Bacillus* subtilisin was 89%. In addition, recombinant *serpin 1270* showed 64% inhibitory activity against trypsin and chymotrypsin (mammalian proteases) and papain (a plant protease). The inhibition profile of purified *serpin 1270* is shown in Fig. 9. Purified recombinant *serpin 1270* inhibited *Bacillus* subtilisin belonging to the S8 family, a subfamily of the MEROPS peptidase database (4).

Serpins are the largest superfamily of protease inhibitors. More than 1500 serpin genes have been identified in animals, plants, poxviruses, bacteria, archea, viruses, and multicellular eukaryotes. Very little is known about the serpin genes of prokaryotes. Serine proteases are the major target of serpins, but a few serpins inhibit other proteases such as cysteine proteases. C. thermocellum can convert cellulose directly into ethanol by its cellulosomal complex, which comprises more than 20 different cellulases and protease inhibitors.

## Conclusion

*C. thermocellum serpin 1270* is active against subtilisin (a bacterial protease) with 89% inhibition and against trypsin and chymotrypsin (serine proteases) and papain (a cysteine protease) with 64% inhibition. SDS-PAGE analysis revealed that the molecular weight of Serpin 1270 was 72kDa. These results demonstrated that serpin 1270 is a potential candidate against protease attack and can be further characterized for the molecular mechanism. The identification of putative serpin 1270 from *Clostridium thermocellum*, which possesses great inhibitory activity against Bacillus subtilisin, provides an insight into purification and characterization of other serpin genes from *C. thermocellum.* This can help us understand the molecular mechanism of Serpin proteins against protease attack in future and further facilitate the activity of cellulases for cellulose degradation in biofuel production.

## Data availability

All the data for this study are contained in the manuscript.

## Acknowledgments

We would like to express our sincere thanks to Prof. Dr. J. H. David Wu (University of Rochester, NY, USA) for his help and kind cooperation during the research.

## Funding and additional information

This work was supported by the Higher Education Commission of Pakistan, and research was performed at the Institute of Biochemistry and Biotechnology, PMAS-Arid Agriculture University, Pakistan under the supervision of Dr. M. Javaid Asad and Prof. Dr. J.H. David Wu at the Department of Chemical Engineering, University of Rochester, Rochester, USA.

## References

1. Rawlings, N., Tolle, D., Barrett, A. (2004) MEROPS: the peptidase database. Nucl. Acid. Res.32, 160–164

2. Law, H., Irving, J., Buckle, A., Ruzyla, K., Buzza, MP., Bashtannyk, T., Beddoe, T. (2005) The high-resolution crystal structure of the human tumor suppressor maspin reveals a novel conformational switch in the G-helix. J. Biol. Chem.280, 22356–22364

3. Bacha, B., Jemel, I., Moubayed, M., Abdelmalek, B. (2017) Purification and characterization of a newly serine protease inhibitor from *Rhamnus frangula* with potential for use as therapeutic drug. BioTech. 7, 148

4. Magalhaes, T., Mambelli, F., Santos, B., Morais, S., Oleveira, S. (2018) Serine protease inhibitors containing a Kunitz domain: their role in modulation of host inflammatory responses and parasite survival. Microbe. Infect. 20, 606–609

5. Viljoen, J., Fred, E., Peterson, W. (1926) The fermentation of cellulose by thermophilic bacteria. J Agri. Sci. 16, 1–17

6. Dzinic, S., Mahdi, Z., Bernardo, M., Vranic, S., Beydoun, H. (2019) Maspin differential expression patterns as a potential marker for targeted screening of esophageal adenocarcinoma/gastroesophageal junction adenocarcinoma. PlosOne. 14

7. Ambadapadi, S., Ramanujam, G., Zheng, D., Sullivan, C., Dai, E., Morshed, S. (2016) Reactive Center Loop (RCL) Peptides Derived from Serpins Display Independent Coagulation and Immune Modulating Activities. J. Biol. Chem. 291, 2874–2887

8. Irving, J., Shushanov, S., Pike, N., Popova, Y., Bromme, D., Coetzer, T. (2002). Inhibitory activity of a heterochromatin-associated serpin (MENT) against papain-like cysteine proteinases affects chromatin structure and blocks cell proliferation. J. Biol Chem. 277, 192–1320

9. Kang, S., Barak, Y., Lamed, R., Bayer, A., Morrison, M. The functional repertoire of prokaryote cellulosomes includes the serpin superfamily of serine proteinase inhibitors. Mol. Microbiol.10, 1365–2958

10. McBee, R. (1954) The characteristics of *Clostridium thermocellum*. J. Bacteriol.67, 505–506

11. Artzi, L., Bayer, E., Morais, S. (2017) Cellulosomes: bacterial nanomachines for dismantling plant polysaccharides. Nature Rev. Microbiol. 15, 83–95.

12. Sanrattana, W., Maas, C. Maat S. (2019) SERPINs—From Trap to Treatment. Front. Med. 6, 25.

13. Laemmli, U.K. Cleavage of structural proteins during the assembly of the head of bacteriophage T4. Nature. 15, 680–685

14. Shrivastava, B., Ghosh, A. Protein purification, cDNA cloning and characterization of a protease inhibitor from the Indian tasar silkworm, *Antheraea mylitta*. Ins. Biochem. Mol. Biol.33, 1025–1033

